# Tn-Seq-based identification of genes that play a role in antimicrobial tolerance of *Pseudomonas aeruginosa* aggregates

**DOI:** 10.64898/2026.11.03.763911

**Authors:** Lise Goltermann, Stefano Gualdi, Morten Rybtke, Shahab Shahryari, Ziwei Liang, Shiyu Lin, Kasper Nørskov Kragh, Leo Eberl, Tim Tolker-Nielsen

## Abstract

Microbial biofilms, whether surface-associated or in the form of aggregated microbes, cause a variety of persistent infections due to their inherent tolerance to antibiotic treatment. Current knowledge of the mechanistic basis of biofilm-associated antibiotic tolerance is fragmented, and the relative importance of previously identified factors remains unknown. We developed a Tn-Seq-based workflow that enables simultaneous assessment of the relative antibiotic tolerance of thousands of individual clonal bacterial aggregates, each established by a single transposon mutant. Using this genome-wide Tn-Seq analysis approach, we identified several factors and pathways that play a role in antimicrobial tolerance o*f Pseudomonas aeruginosa* aggregates. The majority of these factors are important for aggregate-associated tolerance to specific antibiotics, whereas some factors play a role in aggregate-associated tolerance regardless of the class of antibiotic. This methodology provides a framework that can be adapted to investigate aggregate-associated tolerance for other bacterial species and in response to different antimicrobial agents and stressors.

## Introduction

It is well established that bacteria exist not only as solitary single cells, but also in the form of biofilms ^1^. In biofilms, the microbes are located in densely packed microcolonies embedded in a matrix of exopolymers, and in this state the bacteria can tolerate treatment with high doses of antibiotics ^2–4^. Due to their ability to withstand antibiotic treatment, biofilms are a common cause of persistent infections, leading to increased hospitalizations, patient morbidity and reduced quality of life ^1^. It is estimated that about 60–70% of hospital acquired infections are caused by biofilms ^5^. Given that current treatment regimens often fail to eradicate biofilm-based infections, there is an urgent need for novel therapeutic strategies. However, our present knowledge of the mechanistic basis underlying the biofilm-associated antibiotic tolerance remains fragmented and incomplete. Advancing this knowledge is essential for the development of effective new therapies targeting biofilm-related infections.

The recalcitrance of biofilms to antibiotic treatment is commonly attributed to several factors: 1) restricted antibiotic diffusion, particularly in the case of antibiotics that bind to biofilm matrix components, 2) low physiological activity of a subpopulation of the bacteria within the biofilm, 3) expression of specific antibiotic tolerance mechanisms, and 4) the presence of persister cells ^2,3^. The phenomenon is often designated as “biofilm-associated antibiotic tolerance” although it includes both *bona fide* tolerance mechanisms and adaptive resistance mechanisms ^2–4^.

Our current understanding of the underlying mechanisms of biofilm-associated antibiotic tolerance is primarily the result of meticulous studies involving phenotypic characterization of individual mutants ^2,6^. However, the complete set of genes involved in biofilm-associated antibiotic tolerance in bacteria can potentially be identified by the use of next-generation DNA sequencing-based methods such as transposon sequencing (Tn-Seq) ^7^. This is possible because the Tn-Seq method can provide genome-wide maps of bacterial genes that are essential or favorable for survival/proliferation under a particular condition ^8,9^. The Tn-Seq procedure initially involves the generation of a representative mutant library. The library is subsequently propagated under a particular condition, e.g. as a biofilm in the presence of antibiotics. Subsequently, DNA is isolated from the bacteria and PCR amplification of the transposon-gene junction sequences in the mutant pool is performed. By comparison to an untreated control, genes that play a role in survival of biofilm-grown bacteria during antibiotic treatment can then be identified and analyzed to elucidate the molecular mechanisms involved in biofilm-associated antibiotic tolerance. This enables the identification of factors involved in antimicrobial tolerance, by identifying mutants which are depleted during the selection.

Traditionally, biofilms have been grown and studied *in vitro* as surface-attached populations, either statically submerged or under medium flow ^10^. These types of assays, however, may not be well suited for Tn-Seq-based identification of antibiotic tolerance genes, as inter-mutant complementation can obscure the selection of mutants ^11,12^. E.g., Schinner *et al.* employed Tn-Seq on surface-attached populations and found that alginate deficient *P. aeruginosa* mutants benefitted from exploiting alginate overproduction by other Tn mutants within the biofilm while they themselves had a growth benefit by avoiding synthesis of the costly polysaccharide ^11^. An optimal assay for Tn-Seq-based identification of antibiotic tolerance genes should allow each mutant to form an independent biofilm while mimicking the growth mode of biofilms observed in clinical infections. Biofilms found in cystic fibrosis-related pneumonia and chronic wounds are commonly described as small non-attached aggregates suspended in host secretions ^13–15^. Incorporating this non-attached aggregate phenotype into a Tn-Seq assay would therefore enhance the physiological and clinical relevance of the findings, providing a more accurate model for identifying genes involved in biofilm-associated antibiotic tolerance.

The aim of this study was first to develop a Tn-Seq based procedure capable of simultaneously assessing the relative antibiotic tolerance of thousands of individual *P. aeruginosa* aggregates, each initiated by a single transposon mutant; and second to gain mechanistic insights into aggregate-associated antibiotic tolerance in *P. aeruginosa*, a well-established model organism in biofilm research ^16^.

## Results

### Development of a Tn-Seq approach to analyze the genetic basis of aggregate-associated antibiotic tolerance

We developed an experimental setup suitable for Tn-Seq guided identification of genes involved in antibiotic tolerance of *P. aeruginosa* aggregates (Fig. 1). The cultivation was designed to allow individual mutants of a transposon mutant library to form clonal aggregates, thereby reducing inter-mutant complementation. Moreover, the aggregates needed to be grown under conditions that allow antibiotic treatment as well as subsequent removal of the antibiotic. To this end, we inoculated molten LB agar (0.8%) with diluted overnight cultures of wild-type *P. aeruginosa* PA14 and cast the agar as a thin (1 mm) layer in large (Ø 14 cm) petri dishes. The diluted overnight cultures used to inoculate the agar were vortexed thoroughly to ensure that bacteria were evenly dispersed as single cells at the time of inoculation. To avoid extensive bacterial growth and spread at the agar-plastic interface or on the top of the agar, the thin bacteria-containing agar layer was sandwiched between two thin (1 mm) sterile LB agar layers. Upon incubation, aggregates formed within the agar by growth from single immobilized bacteria, as bacterial motility was restricted by the agar matrix ^17^.

**Figure 1.**
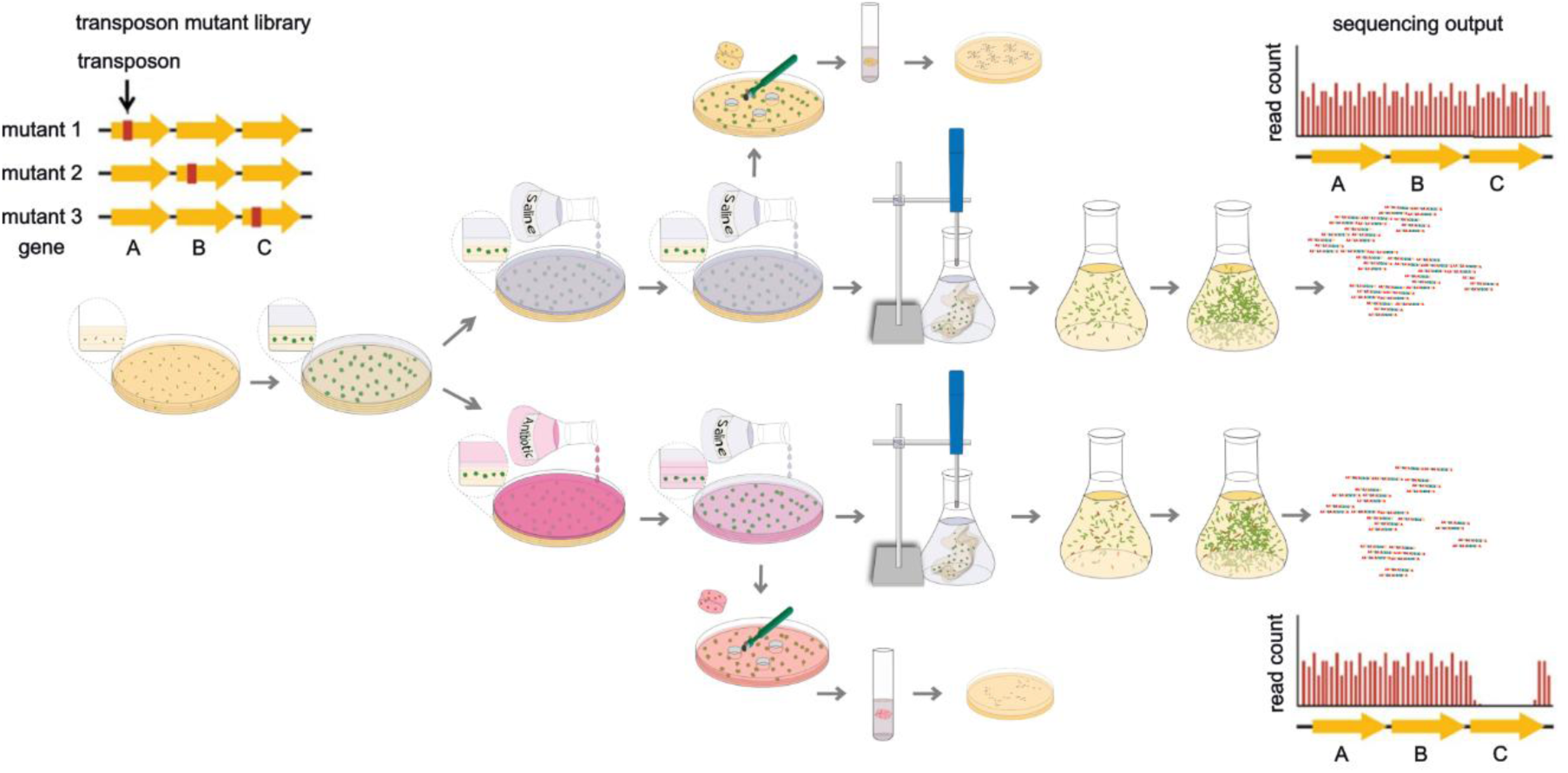
Cartoon illustration of the workflow for Tn-Seq-guided identification of genes that play a role in antimicrobial tolerance of *P. aeruginosa* aggregates. Bacteria from a transposon mutant library are cast in a petri dish in the middle layer of an LB agar sandwich. After 16 hours of incubation, the aggregates are treated for 24 hours either with ciprofloxacin, tobramycin or saline by flooding the plates. At the end of treatment, the antibiotic is removed through several rinsing steps with saline. The bacterial population of each agar plate is subsequently released by homogenization into liquid LB cultures. The surviving cells are allowed to grow for three generations twice, diluting out DNA from dead cells. Subsequently, the number of each of the mutants in the populations is determined by Tn-Seq. In this illustration, disruptions in gene C increases susceptibility and these mutants are counter selected.

In order to assess the growth of bacteria embedded in the sandwich agar plates, samples of the agar gels were harvested from the petri dishes at different time points, and the number of colony forming units (CFUs) was determined by disintegration of the aggregates followed by plate counting. Growth of the wildtype PA14 strain reached stationary phase after 18-20 hours (Fig. 2a). Antibiotic treatments performed in the present study were applied to 16 hour-old aggregates. At this time point, the bacteria in the aggregates were still growing, but their growth rate was declining. We consider these conditions to be clinically relevant, as evidence for limited growth of *P. aeruginosa* in CF lungs has been reported ^14,18^. Each plate was inoculated with approximately 5×10^4^ CFUs (Fig. 2a, time zero). Assuming that each of these bacteria grew into an individual aggregate, the resulting average CFU per aggregate was 3×10^5^ after 16 hours of growth. By enumerating the CFUs in the inoculum and counting the resulting aggregates in a section of the agar sandwich plate, we confirmed that each inoculated bacterium gave rise to an individual aggregate (data not shown). To further verify this, we inoculated plates with a 1:1 mix of CFP-tagged and YFP-tagged *P. aeruginosa* strains as inoculum for aggregate formation and observed that the formed aggregates only displayed a single color of fluorescence, thereby confirming that the aggregates were indeed monoclonal (Fig. S1).

**Figure 2.**
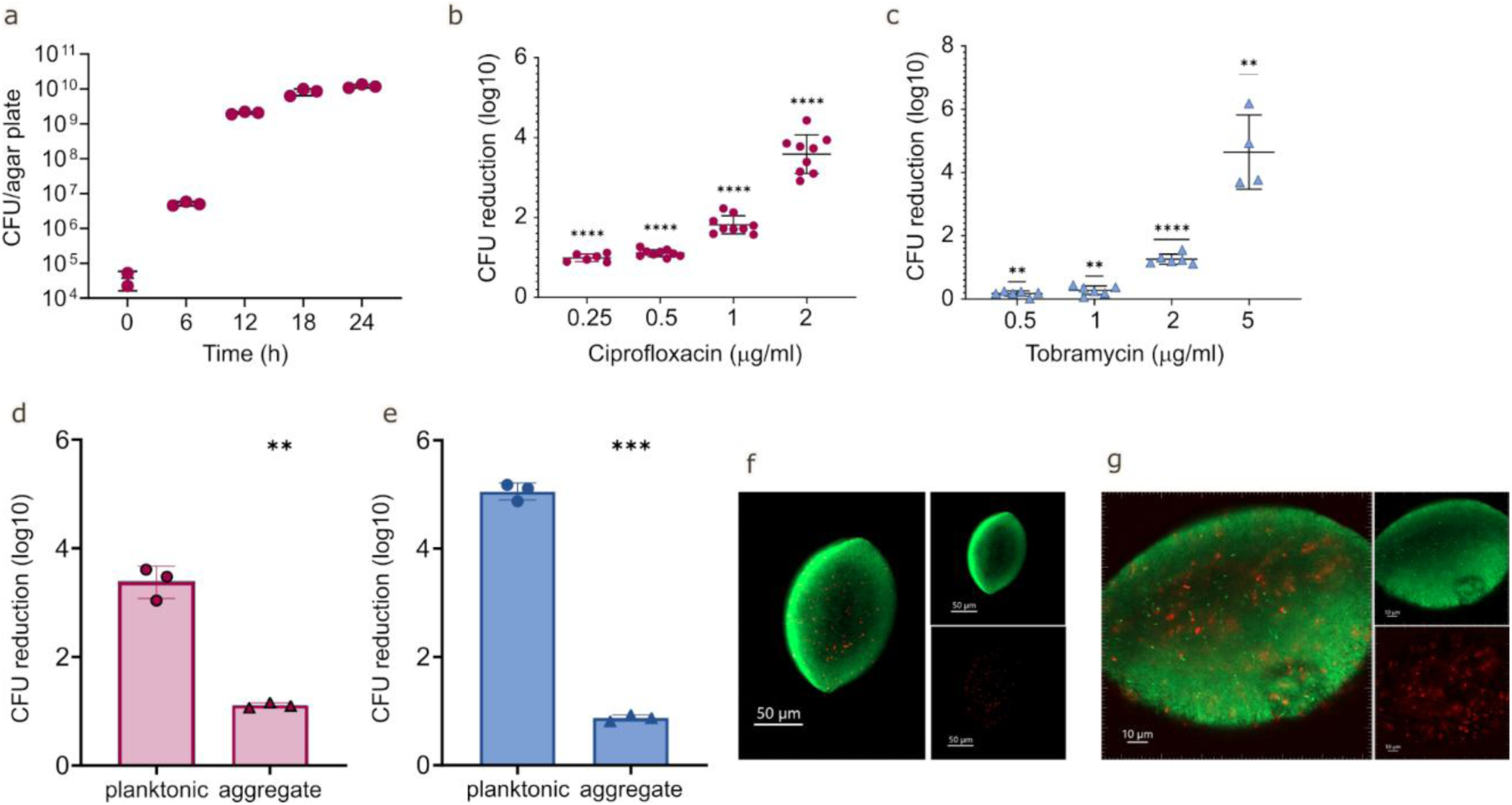
Characterization of the aggregate model. **(a)** Number of colony forming units (CFU) per agar plate of *P. aeruginosa* PA14 aggregates in the sandwich agar plates at increasing incubation time. Mean and standard deviation of 3 biological replicates is shown. **(b** and **c)** The effect of ciprofloxacin **(b)** or tobramycin **(c)** treatment on PA14 agar-embedded aggregates. The aggregates were grown for 16 hours and were then treated for 24 hours with the indicated concentration of antibiotics. Next, the antibiotics were removed by washing and the aggregates were disintegrated, serially diluted and plated for enumeration of viable bacteria (CFUs) and calculation of fold reduction mediated by the antibiotic treatment. Data shows 2-3 biological replicates with 3 technical replicates each, mean and standard deviations are indicated. The significance of the difference of each treatment group from 0 is calculated using a one sample t-test and indicated by asterisks: **p<0.01, **** p<0.0001. (**d** and **e**) Ciprofloxacin **(d)** or tobramycin **(e)** tolerance in planktonic versus aggregate growth. Reduction in CFUs after 24 hours treatment with 0.5 µg/ml ciprofloxacin or 2 µg/ml tobramycin of 16-hour-old aggregates or a similar non-aggregated (planktonic) inoculum. 3 biological replicates, mean and standard deviations are indicated. Student’s t-test ***p<0.001. **(f** and **g)** Confocal laser scanning micrographs of Live/Dead stained 16-hour-old *P. aeruginosa* aggregates. Living cells stain green while red color indicates dying bacteria with compromised membranes or extracellular DNA. Small panels show Cyto9 (green, upper panel) or PI (red, lower panel) signal with the large panel showing the overlay of the two channels. Z-sections are shown here whereas z-stacks are shown as 3D presentations in the supplementary material. The size bars correspond to 50 μm and 10 μm, respectively.

### Characterization of bacteria growing in aggregates

Next, we determined whether the bacteria growing in the aggregates displayed increased antibiotic tolerance compared to planktonically grown cells. We treated 16-hour-old aggregates of PA14 with ciprofloxacin at the concentrations 0.25, 0.5, 1 and 2 μg mL^-1^ and determined the CFUs and log reductions of each treatment compared to a non-treated control sample. Treatment with 0.5 μg mL^-1^ (i.e. 4 x MIC and the concentration used for the mutant library screening) resulted in 1-2 log10 reductions of the number of viable bacteria in the aggregates relative to the untreated control, whereas more pronounced killing required higher ciprofloxacin concentrations (Fig. 2b). Similarly, treatment of the aggregates with tobramycin at different concentrations, showed that 2 µg/ml tobramycin (i.e. 2 x MIC) resulted in approximately a 1 log10 reduction in CFUs (Fig. 2c). For direct comparison of planktonic and aggregated cells under the same conditions, planktonic bacteria were encased in agar with the number of CFUs in the planktonic inoculum matching that of the number of CFUs per agar plate after 16 h of aggregate formation. This ensured that both conditions began with equivalent bacterial loads at the start of antibiotic treatment. Upon immediate treatment of these high-inoculum planktonic cultures with ciprofloxacin or tobramycin, respectively, CFU reductions were observed that were over 100-fold greater than those seen in the aggregates (Fig. 2d and e). These results are consistent with previous work ^17^ demonstrating that the aggregates exhibit increased antimicrobial tolerance compared to their planktonic counterparts, a hallmark of the biofilm mode of growth.

We then assessed whether antibiotic tolerance increased with aggregate age. Aggregates were allowed to form for 16 h, 20 h or 24 h and were subsequently treated with 2 ug mL^-1^ ciprofloxacin for 24 h. Antibiotic susceptibility was determined by comparing CFU reductions to an untreated saline control. While 24 hour-old aggregates showed only 1-2 log reductions in CFUs, 16 hour-old aggregates were approximately 100 times more susceptible under the same condition, indicating that tolerance increased as aggregates entered the stationary phase (Fig. S2). These findings further confirm that 16 h old aggregates were still growing and had not yet reached the stationary phase.

For morphological characterization, the aggregates were stained with a Live/Dead stain and imaged by confocal laser scanning microscopy (CLSM). 16 h old aggregates of PA14 were disc shaped, approximately 50-100 µm in length, and consisted of densely packed bacteria the majority of which were alive (Fig. 2f and 2g and Fig. S3). Importantly, aggregates appeared uniform in size and shape suggesting uniform growth conditions throughout the agar layer.

### Tn-Seq analysis of aggregate-associated antibiotic tolerance

For our Tn-Seq analysis, we used a *P. aeruginosa* PA14 transposon mutant library that was reported to consist of more than 500,000 unique mutants ^19^. In our experiment, we cultivated aliquots of the mutant library in 17 large sandwich-agar plates per treatment condition, which in total contained approximately 300,000 aggregates, each formed by a single transposon mutant. The aggregates were grown for 16 hours and then treated with 0.5 μg/ml ciprofloxacin or 2 µg/ml tobramycin for 24 hours. Following removal of the antibiotic by washing, the aggregates were disintegrated into single bacteria which underwent two consecutive steps of amplification growth in antibiotic-free medium, each for three generations. Parallel experiments were done with saline as a mock control. Genomic DNA was subsequently processed using the Tn-circle method ^20^. The identity and relative number of mutants in the pool of antibiotic-treated and non-treated bacteria were then determined by Tn-Seq analysis. Our analysis indicated that the mutant library contained just below 30,000 mutants (Table 1), which is considerably lower than previously reported values ^19^. Moreover, our Tn-seq analysis showed that antibiotic selection reduced the number of unique mutants roughly by a factor of three for both ciprofloxacin and tobramycin (Table 1). Figure 3 shows a graphical representation of the Tn-Seq analysis outcome. Despite the unexpected low number of transposon mutants, Circos plots ^21^ of the untreated and treated libraries show good coverage of transposon insertions across the genome without any significant bias in transposon distribution (Figure 3).

**Figure 3.**
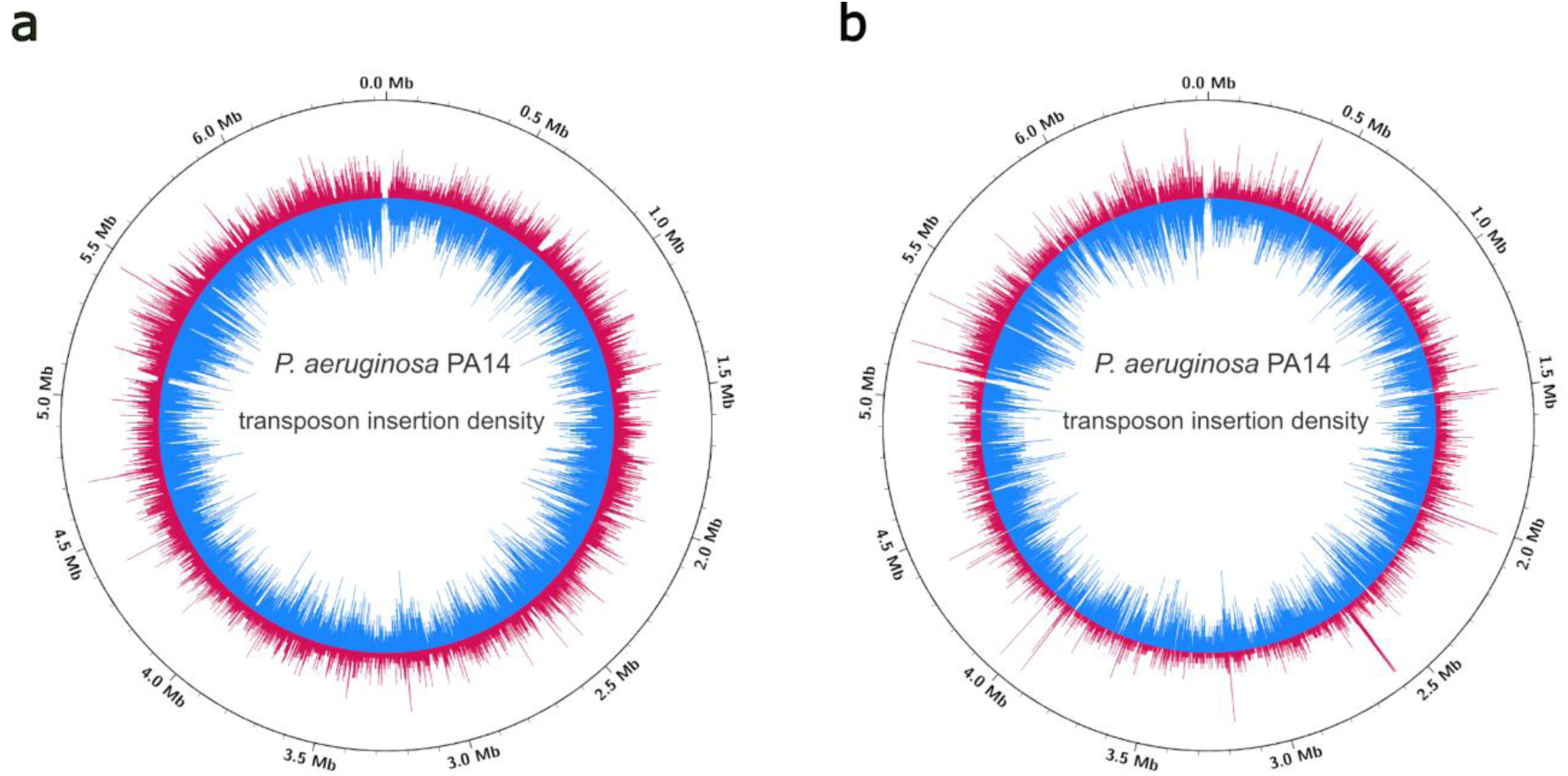
Graphical representation of the outcome of the Tn-Seq-based identification of genes that are involved in antimicrobial tolerance of P. aeruginosa aggregates. Circular plots showing the transposon insertion density in the untreated (blue) and treated (red) samples for ciprofloxacin (**a**) and tobramycin (**b**), respectively. Plots were created using Circos ^21^.

**Table 1.**
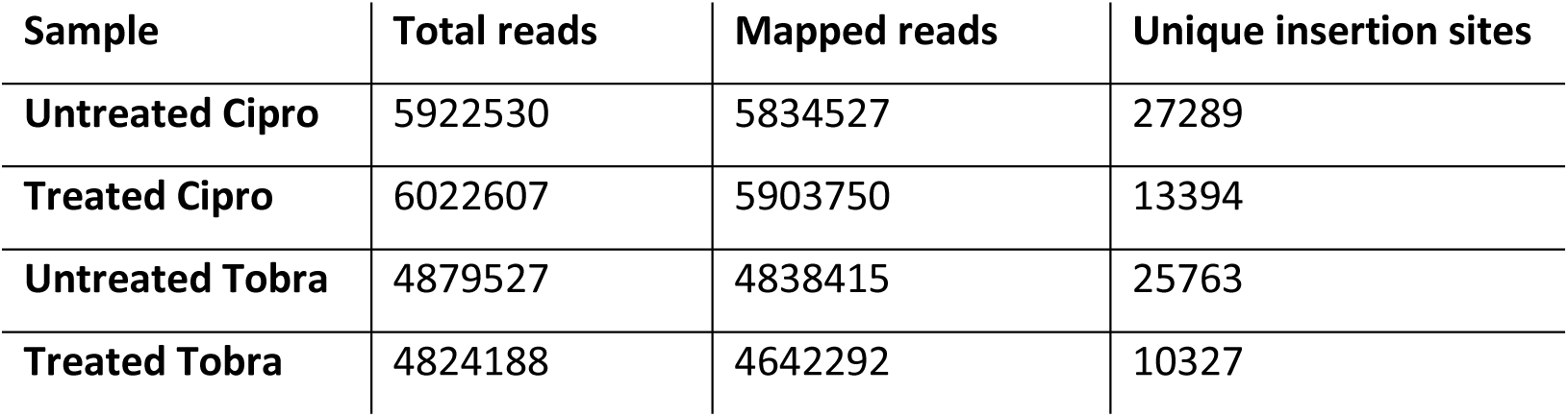
Number of mapped reads and unique transposon insertion sites identified in Tn-Seq for each sample.

| **Sample** | **Total reads** | **Mapped reads** | **Unique insertion sites** |
|---|---|---|---|
| **Untreated Cipro** | 5922530 | 5834527 | 27289 |
| **Treated Cipro** | 6022607 | 5903750 | 13394 |
| **Untreated Tobra** | 4879527 | 4838415 | 25763 |
| **Treated Tobra** | 4824188 | 4642292 | 10327 |

We employed a fitness criterium as defined in the Methods section to rank the relative importance of the antibiotic tolerance genes. The fitness value takes the number of mutants in the initial mutant pool into account, thereby minimizing the influence of random fluctuations of underrepresented mutants. Genes with a low fitness value are characterized by good coverage of transposon insertions and a large difference in abundance between untreated and treated samples, indicating depletion upon antibiotic exposure. Table S2 presents the genome-wide ranking according to the calculated fitness values following ciprofloxacin or tobramycin treatment of the aggregates.

Table 2 and Table 3 present the 10 genes with the lowest fitness values following ciprofloxacin or tobramycin treatment, respectively, indicating a potential role of each of these genes in antibiotic tolerance. Each list includes genes from a variety of functional categories, including genes previously connected with biofilm-associated tolerance, as well as several uncharacterized genes highlighting previously unknown contributors to the tolerance phenotype.

**Table 2.** The 10 genes most important for ciprofloxacin tolerance of *P. aeruginosa* aggregates based on Tn-Seq analysis.

| #Orf | Name | description | PAO1 orthologue | Insertion sites untreated | Insertion sites treated | Total counts untreated | Total counts treated | Fitness |
|---|---|---|---|---|---|---|---|---|
| PA14_23370 | *orfK* | putative UDP-N-acetylglucosamine 2-epimerase | PA3148 | 7 | 0 | 1225 | 0 | -8582 |
| PA14_70120 | - | putative MFS transporter | PA5311 | 9 | 0 | 816 | 0 | -7353 |
| PA14_50840 | - | Putative DNA helicase | PA1045 | 12 | 1 | 1733 | 2 | -6358 |
| PA14_64080 | dipZ | thiol:disulfide interchange protein | PA4845 | 10 | 1 | 1186 | 1 | -5341,5 |
| PA14_62200 | *mrcB* | penicillin-binding protein 1B | PA4700 | 16 | 1 | 3561 | 11 | -4452,5 |
| PA14_69500 | *argH* | argininosuccinate lyase | PA5263 | 7 | 2 | 2358 | 2 | -3931,7 |
| PA14_62910 | *carB* | carbamoylphosphate synthetase large subunit | PA4756 | 16 | 2 | 1006 | 3 | -3524,5 |
| PA14_38730 | *yhhS* | putative MFS transporter | PA1993 | 7 | 0 | 426 | 0 | -2989 |
| PA14_67400 | - | putative ABC transporter periplasmic substrate-binding protein | PA5103 | 5 | 0 | 596 | 0 | -2985 |
| PA14_06600 | - | putative acyl-CoA dehydrogenase | PA0506 | 8 | 3 | 2363 | 3 | -2955 |

**Table 3.** The 10 genes most important for tobramycin tolerance of *P. aeruginosa* aggregates based on Tn-Seq analysis.

| #Orf | Name | description | PAO1 orthologue | Insertion sites untreated | Insertion sites treated | Total counts untreated | Total counts treated | Fitness |
|---|---|---|---|---|---|---|---|---|
| PA14_52260 | *lemA (gacS)* | sensor/response regulator hybrid | PA0928 | 19 | 4 | 12816 | 13 | -13732,5 |
| PA14_15780 | *nagE* | putative Phosphotransferase system N-acetylglucosamine-specific enzyme IIABC | PA3761 | 13 | 0 | 908 | 0 | -11817 |
| PA14_62910 | *carB* | carbamoylphosphate synthetase large subunit | PA4756 | 13 | 0 | 907 | 0 | -11804 |
| PA14_40260 | - | conserved hypothetical protein | PA1874 | 13 | 0 | 793 | 0 | -10322 |
| PA14_52240 | *pirS* | two-component sensor | PA0930 | 12 | 0 | 711 | 0 | -8544 |
| PA14_32390 | *mexF* | RND multidrug efflux transporter MexF | PA2494 | 11 | 0 | 756 | 0 | -8327 |
| PA14_29230 | - | putative hydrolase | PA2699 | 9 | 0 | 903 | 0 | -8136 |
| PA14_30650 | *gacA* | response regulator GacA | PA2586 | 7 | 0 | 1145 | 0 | -8022 |
| PA14_28310 | - | putative enoyl-CoA hydratase | PA2767 | 9 | 0 | 847 | 0 | -7632 |
| PA14_30180 | *idh* | monomeric isocitrate dehydrogenase | PA2624 | 9 | 0 | 833 | 0 | -7506 |

### Aggregate-associated antibiotic tolerance of selected mutants

For the 10 genes with the lowest fitness value from either the ciprofloxacin selection or the tobramycin selection, insertion plots were constructed to visualize the selective depletion of mutants (Figure 4). The insertions plots include each of the genes as well as the flanking upstream and downstream genomic regions. Furthermore, to confirm antibiotic specific effects, the insertions plots from the genes identified in the ciprofloxacin selection were compared to their counterparts from the tobramycin selection and vice versa (Figure S4). The insertion plots clearly show the selective depletion of the mutants by the antibiotic treatments, and at the same time provide information about the number of unique mutants in the mutant library.

**Figure 4.**
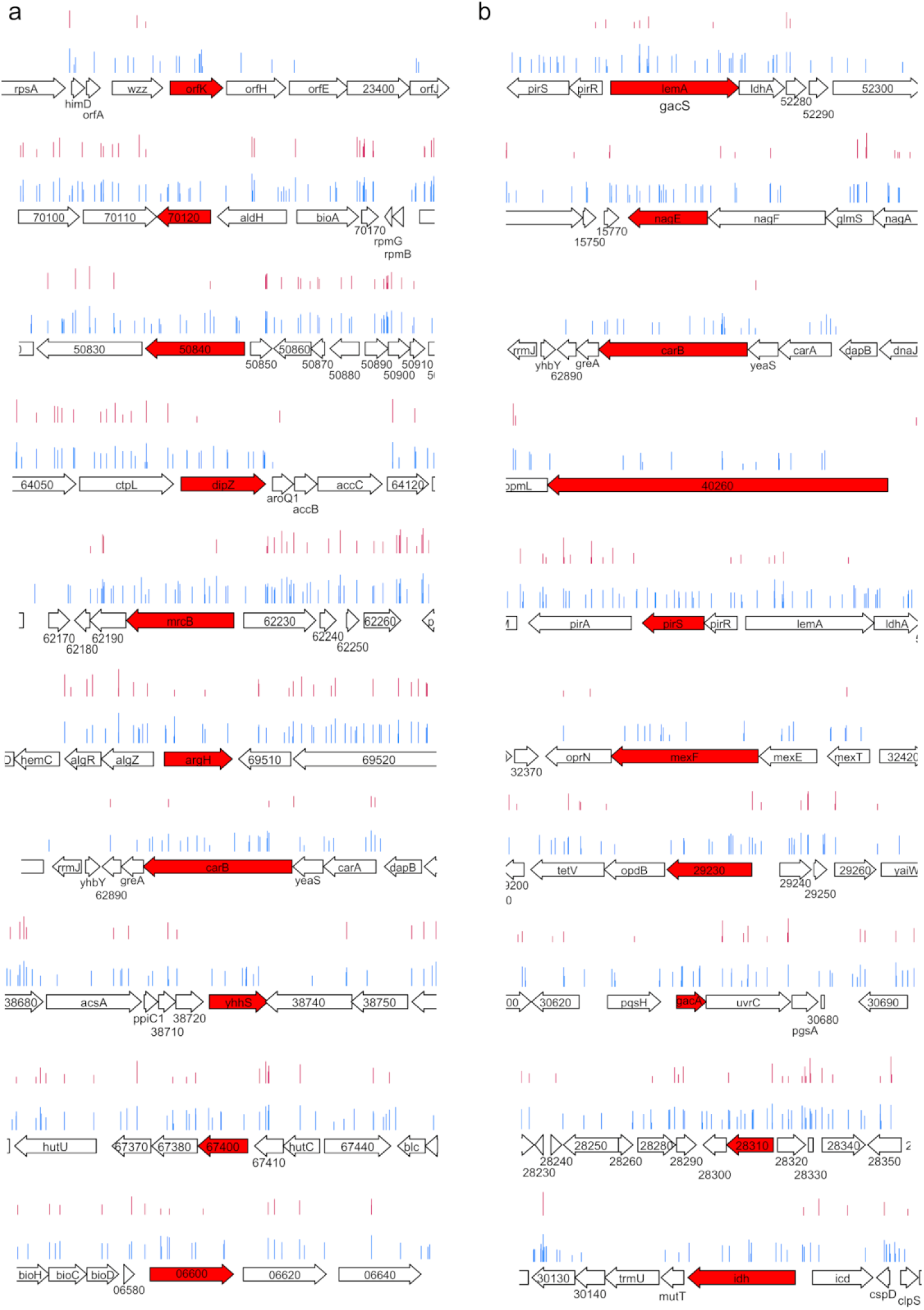
Insertion plots of the genes and flanking regions for the 10 genes with the lowest fitness value from either the ciprofloxacin selection (**a**) or the tobramycin selection (**b**). The location and number of reads for each insertion is shown for the untreated sample (blue) and treated sample (red). The height of each bar signifies the number of counts, plotted as log10(10x) with x being the raw counts.

To validate the Tn-Seq data, we used the fitness-based ranking (Table 2 and 3) along with the insertion plots (Figure 4) to guide the selection of a set of mutants from the independently generated *P. aeruginosa* UCBPP-PA14 mutant collection ^22^ to be tested for alterations in antimicrobial tolerance compared to wildtype PA14. The mutants were disrupted in previously characterized antibiotic tolerance genes as well as in genes with no prior reported association with antibiotic tolerance.

From the ciprofloxacin selection, we tested PA14_70120 encoding a predicted MFS transporter, PA14_62200 encoding MrcB, PA14_62910 encoding CarB, and PA14_06600 involved in fatty acid metabolism. From the tobramycin selection, we tested PA14_52260 encoding GacS (previously designated LemA), PA14_32390 encoding MexF, PA14_30650 encoding GacA, and PA14_62910 encoding CarB. PA14_62910 (*carB*) was the only gene present in the top 10 hits of both screens. All selected transposon mutants were PCR verified and tested in conventional planktonic MIC assays. The PA14_32390 transposon mutant could not be verified by PCR, and therefore a PA14Δ32390 in frame deletion mutant was constructed as described in the Methods section. All strains displayed a ciprofloxacin MIC of 0.125 μg/ml and all but one mutant showed a tobramycin MIC of 1 μg/ml similar to that of the wildtype PA14 (Table S3). Only the PA14_06600 mutant exhibited a 2-fold lower MIC of tobramycin despite being identified in the ciprofloxacin screen. In addition, maximal growth rates measured during exponential planktonic growth were similar across mutants, thus ruling out major growth defects as a cause of altered aggregate-associated tolerance. Only the *carB* mutant showed a slight decrease in maximum growth rate (Table S3).

Each mutant was grown as monoclonal aggregates in the plate model and the tolerance towards treatment with 0.5 μg/ml ciprofloxacin or 4 μg/ml tobramycin was assayed by comparing the CFU reduction upon treatment with that of wildtype PA14 (Fig. 5a-d). All tested strains formed aggregates of similar size based on microscopic inspection (data not shown). Aggregates treated with the saline control showed approximately equal numbers of viable cells after 24 h (Fig. S5), except for the *carB* aggregates that contained about half the number of viable cells compared with the wildtype aggregates. Aggregate-associated antibiotic tolerance was assayed for both ciprofloxacin and tobramycin in all mutants to determine whether susceptibility-changes were antibiotic-specific.

**Figure 5.**
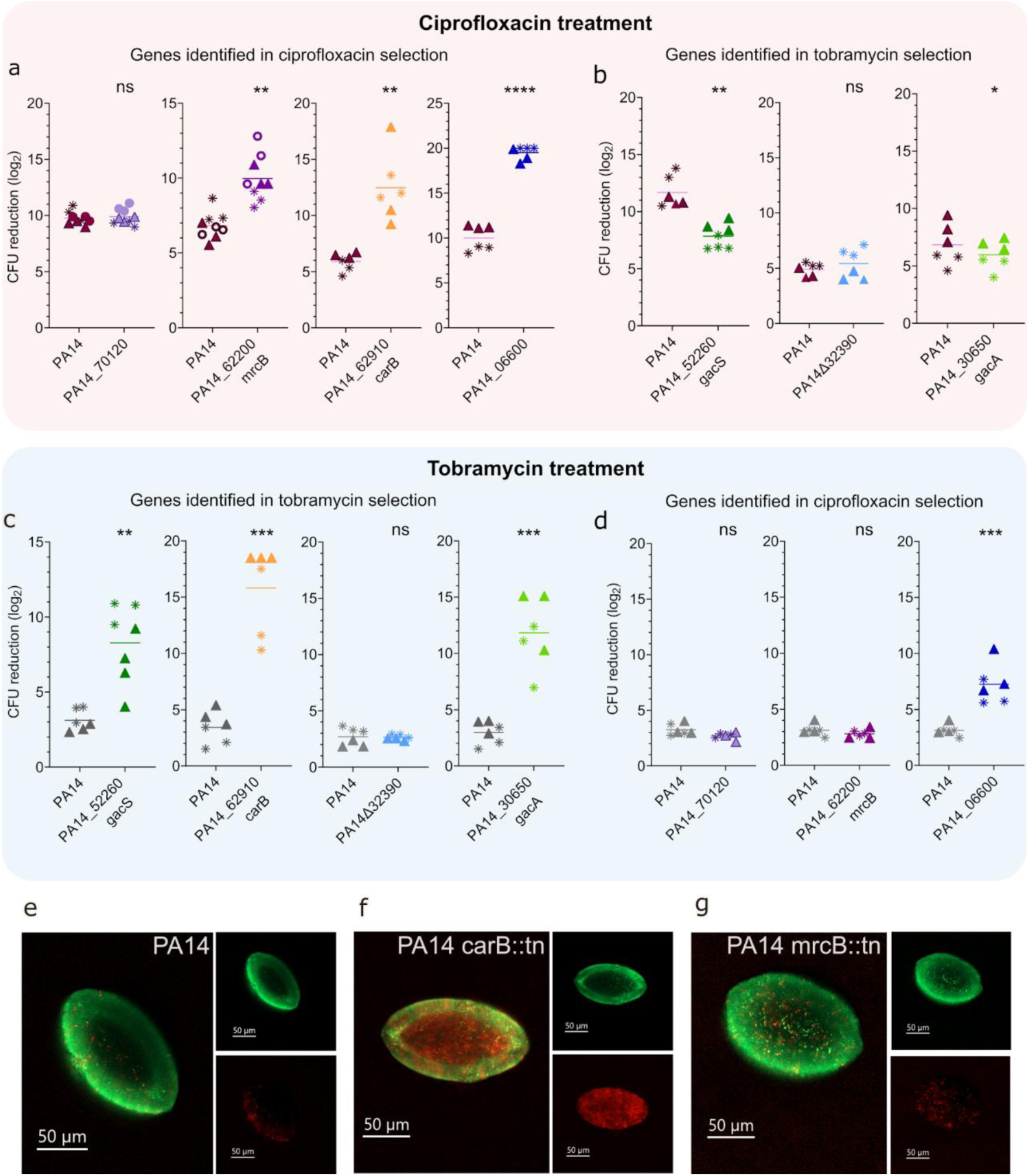
(**a-d**) Antibiotic tolerance of aggregates formed by the indicated *P. aeruginosa* PA14 transposon/deletion mutants and the wild-type PA14. The respective *P. aeruginosa* strains were grown as agar-embedded aggregates for 16 hours. The aggregates were then treated for 24 hours with ciprofloxacin **(a** and **b)** or tobramycin **(c** and **d)**. Next, the antibiotic was removed by washing and the aggregates were disintegrated, serially diluted and plated for enumeration of viable bacteria (CFUs) and calculation of the reduction of CFUs mediated by the antibiotic treatment. The presented data consists of three technical replicates and two biological replicates (indicated by stars and triangles, respectively). Means are indicated by lines. For statistical analysis, each mutant was compared to the wildtype originating only from the same experiment and data compared using Student’s t-test * p<0.05; ** p<0.01; **** p<0.001; ns, not significant. **(e-g)** Confocal laser scanning micrographs of Live/Dead stained 16-hour-old *P. aeruginosa* aggregates of the wildtype (**e**), *carB* mutant (**f**) and *mrcB* mutant (**g**). Living cells stain green while red indicates dead bacteria with compromised membranes. Small panels show Cyto9 (green, upper panel) or PI (red, lower panel) signal with the large panel showing the overlay of the two channels. The size bars correspond to 50 μm.

Three of the four ciprofloxacin-screen-derived mutants (PA14_62200, PA14_62910 and PA14_06600) were significantly more susceptible to ciprofloxacin than the wildtype PA14 in the aggregate assay (Figure 5a). One mutant, PA14_70120, did not show any difference in aggregate-associated ciprofloxacin tolerance when compared with the wildtype. However, this mutant exhibited a delay in post-treatment regrowth in liquid culture relative to the wildtype (Fig. S6). This delay could reflect either a slight reduction in CFU numbers or a delayed onset of growth following treatment. In either case, such effects would be expected to contribute to the depletion of this transposon mutant from the ciprofloxacin-treated library.

Apart from *carB*, none of the mutants selected from the tobramycin screen showed any increase in ciprofloxacin susceptibility (Figure 5b). Interestingly, the *gacS* and *gacA* mutants were slightly less susceptible towards ciprofloxacin compared with the wildtype (Figure 5b)

Three of the four tobramycin-screen-derived mutants (PA14_52260, PA14_62910 and PA14_30650) were significantly more susceptible to tobramycin than the wildtype PA14 in the aggregate assay (Figure 5c). Among these mutants were both members of the GacA/GacS-system, encoded by PA14_30650 and PA14_52260, respectively. This two-component system is part of a complex regulatory network of genes and regulatory small RNAs that control virulence and biofilm formation partly in response to external stressors such as antibiotics ^23^. The *gacA* and *gacS* transposon mutants showed significantly increased susceptibility towards tobramycin (Figure 5c) but not ciprofloxacin (Figure 5b) when growing in aggregates, while MIC values remained unchanged for both antibiotics.

As the only gene identified in the top 10 in both screens, aggregates of the *carB* transposon mutant showed substantially increased susceptibility towards tobramycin compared with the wildtype (Figure 5c).

The *mexF* deletion mutant (constructed because the *mexF* transposon mutant could not be PCR-verified) did not display any changes in susceptibility towards tobramycin or ciprofloxacin (Fig 5b and 5c). Post-treatment re-growth in liquid culture was also unaffected (data not shown). While the in-frame *mexF* deletion mutant showed no phenotype, previous studies have reported altered tobramycin susceptibility for colony biofilms of *mexF* transposon mutants ^24^. Importantly, this effect was attributed to membrane stress caused by truncated MexF variants incorporated into the membrane. This may explain why the in frame *mexF* deletion behaves differently as no truncated protein product is expressed. To confirm this hypothesis, we acquired a previously described *mexF* transposon mutant in PAO1 ^25^. Compared to its respective wildtype, aggregates of this *mexF* mutant were significantly more susceptible to tobramycin while no change in planktonic MIC was observed (Fig. S7 and Table S3) thus confirming our Tn-Seq result.

Interestingly, one of the mutants from the ciprofloxacin selection, PA14_06600, was also susceptible towards tobramycin (Figure 5d), which is consistent with its reduced MIC for tobramycin relative to the wildtype. Closer inspection of the Tn-Seq data revealed that PA14_06600 was also counter-selected under tobramycin treatment (Table S2). PA14_06600 encodes one of several acyl-CoA dehydrogenases in *P. aeruginosa* which may contribute to aggregate maintenance during nutrient limited conditions through β-oxidation of fatty acids and the production of acyl-CoA for central metabolism ^26^.

Since the *carB* mutant was strongly counter-selected during both ciprofloxacin and tobramycin treatment we extended our investigations for this mutant. The *carB* gene (PA14_62910), encoding the large subunit of the carbamoyl-phosphate synthase (CPSase), is in operon with four other genes involved in the biosynthesis of pyrimidine and arginine, which is known to be important for survival under anoxic conditions. Because the *carB* mutant aggregates were highly susceptible to both ciprofloxacin (Fig. 5a) and tobramycin (Fig. 5c), we speculated that these aggregates may be more sensitive to oxygen limitation within the aggregate interior, leading to altered physiology compared with the PA14 wildtype. Using CLSM (Fig. 5 e-g), we analyzed the structural and physiological properties of *carB* mutant aggregates prior to treatment. Live/Dead staining revealed reduced viability in the aggregate interior, while aggregate size and shape remained largely unchanged (Fig. 5f). This aligns well with the viability data shown in Figure S4. In contrast, the *mrcB* transposon mutant (Fig. 5g) formed aggregates indistinguishable from the wildtype (Fig 5e), confirming a different and ciprofloxacin-specific mechanism of tolerance.

As described above, the results of the Tn-Seq experiments were validated by assessing the aggregate-associated antibiotic tolerance of PA14 transposon mutants in comparison to the isogenic wildtype. The fact that independent mutants were counter selected both in the Tn-Seq experiment and in the subsequent verification experiments indicate causality. However, to investigate for polar effects we complemented the transposon mutants that showed a tolerance phenotype. In the case of *mrcB* and PA14_06600, aggregates of the complemented mutants were significantly more antibiotic tolerant than the mutant aggregates (Fig. S8), confirming the role of these genes in antibiotic tolerance. However, for the *carB* mutant we did not see complementation (Fig. S8). The complementation construct consisted of the P_BAD_ promoter fused to the *carB* gene and its upstream region containing a ribosome binding site (as described in the supplementary material). As shown in figure 4, the entire five-gene operon containing *carB* was counter selected by the antibiotic treatments. Our Tn-Seq data and *carB* complementation experiments thus support a role of the five-gene operon (hereafter referred to as the Car operon) in antibiotic tolerance instead of just the *carB* gene. We did not complement the *gacA* and *gacS* mutants, as these genes are positioned far away from each other on the chromosome and validate each other.

## Discussion

Here, we developed a Tn-Seq based procedure that enables simultaneous assessment of the relative antibiotic tolerance of thousands of individual bacterial aggregates. Our model for cultivation of aggregates enables individual mutants from a transposon mutant library to form separate aggregates and thereby reduces inter-mutant complementation that could affect the outcome of the Tn-Seq experiments. Cultivating aggregates in thin agar layers was intended to ensure that all the aggregates were exposed to similar chemical and environmental conditions. Mutants with major aggregate growth defects are minimally represented in the untreated samples, and as a result, our fitness ranking provides an overview of the mutants which are well represented and able to grow as aggregates under untreated conditions but are selectively eliminated from the treated sample.

As expected, bacteria within the aggregates exhibited greater tolerance to antibiotic treatment than planktonic bacteria, which is characteristic of bacteria in the biofilm mode of growth ^2,3^, and corresponds well with previous work on aggregate growth of *P. aeruginosa* ^17^. The size of the aggregates resemble that of *P. aeruginosa* aggregates found in tissue from people with cystic fibrosis-related chronic pneumonia and chronic wounds, where the aggregates are often found to be in the range of 5-100 µm in diameter ^13–15,27^. While our model appears to reflect clinically relevant biofilm structures, future studies may benefit from optimizing the growth medium composition to further enhance its physiological relevance ^28^.

One important limitation of this study is the relatively low density of the transposon mutant library. A higher density transposon mutant library would greatly increase the resolution of the gene ranking. However, despite the mutant library not being as dense as expected, we identified several genes that were demonstrated to play a role in aggregate-associated tolerance.

The results were validated through direct comparison of aggregate-associated antimicrobial tolerance between selected mutants and the wildtype PA14, as well as by comparison with other studies. We identified both members of the GacS/GacA system in the tobramycin screening. Activation of the Gac-system leads to biofilm maturation and production of the pel and psl matrix components which are known to play a role in tolerance to tobramycin ^29^. This is well in line with the observation that mutations in the Gac-system were counter selected during tobramycin but not ciprofloxacin treatment. Importantly, Gac mutants in *Pseudomonas* species have been found to co-exist with the wild-type as both cheaters and mutualists thus validating our approach in identifying mutants which may not be depleted from a mixed population ^30^

Unexpectedly, both the GacS and GacA transposon mutant showed a slight increase in aggregate-associated ciprofloxacin tolerance. While the mechanism underlying this effect remains unclear, the Gac-system regulates a wide array of cellular functions including quorum sensing, secondary metabolism, motility and secretion systems, any of which may influence the response to ciprofloxacin ^23^.

We also found transposon mutants of PA14_40260 and PA14_52250 (*pirS*) to be counter-selected under tobramycin treatment. The PAO1 ortholog of PA14_40260, termed PA1874, has been characterized as part of an efflux pump that is expressed during biofilm growth resulting in tolerance to tobramycin ^31^. PirS positively controls the porin OprH which is known to play a role in aminoglycoside susceptibility ^32^.

One gene, *carB*, ranked among the top ten in both antibiotic screens. We speculate that this result reflects the importance of arginine metabolism under oxygen-limited conditions, such as those likely present in the interior of the aggregates. In surface attached biofilms, antibiotic tolerant subpopulations are often located in deeper layers of the biofilm ^33,34^, a microenvironment that resembles the interior of aggregates. In the case of the *carB* mutant, the viability of this subpopulation appears to be greatly reduced as suggested by CFU count and CLSM imaging. This may explain the increased antibiotic susceptibility observed in this strain and suggests that metabolic adaptation to oxygen limitation plays a critical role in sustaining antibiotic-tolerant subpopulations within aggregates ^35^. This aligns well with previous work, identifying amino acid transporters as important contributors to the aggregate phenotype ^36,37^. Notably, *carB* has been implicated in persister cell formation in planktonic *P. aeruginosa* cultures ^38^, likely due to the Car operon’s central role in both pyrimidine and arginine biosynthesis, linking it to multiple essential metabolic pathways.

Katharios-Lanwermeyer and O’Toole (2022) have suggested that maintenance of a persistent biofilm is an active process ^39^. They emphasized that energy-conserving pathways, such as arginine fermentation, may play a key role in the maintenance of these communities. Furthermore, targeting maintenance pathways was suggested as a novel strategy for the treatment of biofilm-associated infections. These considerations agree well with our findings of a role of the Car operon in antibiotic tolerance of *P. aeruginosa* aggregates.

Mutants of *carB*, and *mrcB* have previously been shown to exhibit increased susceptibility to ciprofloxacin in a colony-biofilm model ^40^. Several genes with low fitness values in our study (PA14_23370 *orfK*, PA14_57650 and PA14_20030 *hasD*) have previously been linked to aggregate or biofilm growth and antimicrobial tolerance ^36,41–43^. Disruption of these genes may therefore result in an aggregate phenotype with increased susceptibility to one or more antibiotic classes. The mutants displaying the strongest phenotypes may be specifically impaired in supporting the survival and proliferation of antibiotic tolerant cells within the aggregate interior.

The concept that biofilm-associated antibiotic tolerance is multifactorial is well established. However, to our knowledge, this is the first study to provide experimental evidence based on the simultaneous assessment of antibiotic tolerance across thousands of individual biofilm aggregates. Our findings align with work by Stewart et al (2019), who used transcriptomic, metabolomic, physiological, and computational modeling approaches to investigate the mechanisms of antibiotic tolerance in *P. aeruginosa* biofilms grown in drip flow reactors ^44^. Their results, along with other studies ^40,45,46^, support the view that antibiotic tolerance arises from an interplay of factors including limited antibiotic diffusion, altered metabolism, differential gene expression, and specific physiological adaptations ^44^.

In conclusion, we have established a Tn-Seq based procedure that enables simultaneous evaluation of antibiotic tolerance across thousands of individual bacterial aggregates. By applying this genome-wide Tn-Seq analysis approach, we provided evidence that antimicrobial tolerance of *P. aeruginosa* aggregates depends on a broad array of genes spanning multiple functional categories. These findings underscore the inherently multifactorial nature of biofilm-associated antibiotic tolerance, which arises from the combined effect of numerous, functionally diverse genetic determinants.

## Methods

### Bacterial strains and growth media

All bacterial strains and primers used in this study are listed in Table 4 and Table S1, respectively. Defined transposon mutants were verified by PCR. The transposon mutant library was constructed in *P. aeruginosa* PA14 by Vitale et al. ^19^. The PA14Δ32390 in frame deletion mutant was made according to Hmelo et al. ^47^. *P. aeruginosa* PA14 was grown in Lysogeny Broth (LB) at 37°C with shaking for generation of planktonic overnight cultures. LB solidified with 2% agar was used for determination of colony forming units (CFUs). LB solidified with 0.8% noble agar (Fisher Scientific, Denmark) was used for cultivation at 37°C of bacteria as agar-embedded aggregates.

**Table 4.** Bacterial strains used in the study.

| ***P. aeruginosa* strains** | **Description** | **Reference** |
|---|---|---|
| PA14 (WT) | Wild type PA14 | <sup>22</sup> |
| PA14_70120 | PA14_70120 transposon mutant | <sup>22</sup> |
| PA14_62200 | PA14_62200 transposon mutant | <sup>22</sup> |
| PA14_06600 | PA14_06600 transposon mutant | <sup>22</sup> |
| PA14_62910 | PA14_62910 transposon mutant | <sup>22</sup> |
| PA14_52260 H9 | PA14_52260 transposon mutant | <sup>22</sup> |
| PA14_30650 F7 | PA14_30650 transposon mutant | <sup>22</sup> |
| PA14Δ32390 | In frame deletion mutant | This study |
| PAO1 | Wild type PA01 | Lab collection |
| PAO1-YFP | PAO1 tagged with eYFP in a mini-Tn*7* construct, Gm<sup>r</sup> | <sup>51</sup> |
| PAO1-CFP | PAO1 tagged with eCFP in a mini-Tn*7* construct, Gm<sup>r</sup> | <sup>51</sup> |
| *mexF*::tn52 | Transposon mutant in *mexF* of PAO1 | <sup>25</sup> |

### Determination of minimum inhibitory concentration (MIC) values

Minimum inhibitory concentration (MIC) values were determined by broth microdilution according to standard protocols in LB medium. Briefly, 190 μl bacterial cell culture containing ≈10^5^ cfu/ml was dispensed into a 96-well plate along with 10 μl antibiotic. The plate was incubated in a Tecan Sunrise plate reader at 37°C for 20 h, and OD was measured every 20 min at 595 nm. The MIC was determined as the lowest concentration, which inhibited visible growth (OD < 0.1) in the wells. Maximum growth rates were calculated from the growth curves of untreated cultures upon the entry into logarithmic phase by fitting 5-7 datapoints to the Exponential Growth Equation in GraphPad Prism 10.6.1. All experiments were conducted in minimum 3 biological replicas.

### Cultivation of P. aeruginosa aggregates in sandwich agar plates

*P. aeruginosa* aggregates were grown in a three-layered LB agar sandwich cast in Ø14 cm Petri dishes. The model included three layers in order to trap individual cells only in the middle layer, resulting in non-attached aggregates forming in this layer, and preventing surface growth on the dish itself or the agar’s top surface. Thus, a homogeneous distribution of aggregates could be created in one layer in an effort to let all aggregates experience identical environmental conditions. The first 1 mm thick agar layer was created by pouring 14.4 mL sterile molten LB agar (48-50°C) into a Ø14 cm Petri dish which was left to harden for 30 min. Then a second 1 mm thick agar layer with bacteria was created by adding 14.4 mL of LB-agar containing *P. aeruginosa* bacteria diluted to an approximate final concentration of OD_600_ 1 x 10^-6^. The second layer was left to harden for 30 min. A final 1 mm agar layer was cast on top of the second layer by adding 14.4 mL sterile LB-agar that was left to harden for 30 minutes. The finished sandwich agar plates were incubated upside-down at 37°C until determination of colony forming units (CFU) or antibiotic treatment was performed.

High-inoculum planktonic plates were made using over-night stationary phase cultures. To prepare the inoculum, bacterial cells were harvested and resuspended in LB broth to an OD600 of 10 corresponding to ≈10^10^ cfu/ml. Subsequently, 250 μl was inoculated into the 6.4 ml molten 0.8% LB agar used to form the bacteria-containing middle layer.

### Determination of colony forming units (CFUs)

Antibiotic mediated killing was assessed based on CFU count. After antibiotic treatment (see below), three samples were isolated from each plate using a 10 mm punch biopsy knife to release an agar plug containing aggregates. Each plug was combined with 950 µl saline and 2 ceramic beads in a 2 ml screw cap tube and homogenized for 2 times 10 seconds in a MagNALyser (Roche). Aggregates were disintegrated by 5 min degassing followed by 5 min sonication in an ultrasonication bath. Samples were then serial diluted and plated on LB agar plates for subsequent CFU count. In parallel to the CFU determinations, each dilution was tested for its ability to regrow. For this, 10 µl of each dilution was added to 190 µl LB in a 96-well plate and incubated at 37°C with shaking in a Tecan Sunrise plate reader with OD600 measured every 20 min for 20h.

### Antibiotic treatment of P. aeruginosa aggregates

For the Tn-Seq experiments, we employed a previously constructed and characterized Tn5 transposon mutant library in *P. aeruginosa* PA14 ^19^. The transposon mutant library stored at −80°C was thawed and bacteria were diluted according to the OD values and cased into the sandwich aggregate plate model as described above. For antibiotic treatment of the aggregates in the sandwich agar plates, 43,2 mL saline (0.9% w/v NaCl) containing the double antibiotic concentration of the desired final concentration was added on top of the sandwich agar plates after 16 hours of incubation at 37°C. Subsequently, the final antibiotic concentration was reached as the antibiotic diffused into the agar. Control plates were treated with 43,2 mL saline. Plates were incubated with shaking at 40 rpm for 24 hours with antibiotic treatment or saline at 37°C. After 24 hours, the excess fluid on top of the agar was removed, and in order to remove the antibiotic, the plates were washed three times with saline at 37°C with shaking at 40 rpm for 30 min to let antibiotics diffuse out of the agar. Control plates were rinsed as well.

### Preparation of aggregate populations for DNA extraction

The bacteria bound in the aggregates in the plates were released by homogenization. Agar gels were released from the plate using a sterile cell scraper and placed in an Erlenmeyer flask (4 plates per flask) and the volume brough to 400ml with LB. The gels were homogenized for 1 min at 16,000 rpm and the flasks were sonicated (5 min degas, 10 min sonication) to separate the cells from agar remnants. Released bacteria were transferred into a 2L conical flask and the volume brought to 1L with pre-warmed LB broth and underwent amplification-growth until reaching an OD_600_ of 0.7 at 200 rpm at 37°C. As the cultures reached OD_600_ 0.7, the cultures were diluted 20 times in 1L fresh pre-warmed LB broth and regrown to OD_600_ 0.7 under the same conditions. Samples were harvested as 30 ml aliquotes which were pelleted (8000rpm, 10 min) and stored at −20°C until DNA extraction. The number of CFUs/ml was sampled at each stage of re-growth to ensure library coverage and identify possible bottlenecks.

### DNA extraction, sequencing of the libraries, data analysis and bioinformatics

DNA was extracted from the samples using the Qiagen Blood&Tissue Kit according to the manufactures guide for Gram-negative bacteria. The DNA was subsequently processed using the Tn-circle method ^20^ with few modifications. Briefly, the genomic DNA was sheared into 300 bp long fragments using Covaris. NebNext Ultra II DNA library Prep Kit (Cat. No. E7645S) was employed for end repair and adaptor ligation. Size selection of DNA fragments was performed on 1.5% agarose gels and recovered with QIAquick Gel Extraction Kit (Qiagen Cat. No.28704). Following the circularization, DNA was amplified using Phusion High-Fidelity DNA polymerase (Thermo Fisher Scientific, Cat. No. F530L) and the primers shown in Table S3. The libraries were sequenced using MiSeq reagent kit (V2, 300 cycles; Illumina). The FASTQ reads were demultiplexed and trimmed using the command line “cutadapt -a adapter -q output.fastq.gz ^48^. The reads were mapped onto the genome of *P. aeruginosa* UCBPP-PA14 using the software Tn-Seq Explorer and TRANSIT ^49,50^. The unique insertion count (UIC) specifies how many times a transposon was found in the coding region of the gene (discarding the 5% at the beginning and 20% at the end of the ORF). To enable downstream data analysis, genes with an UID of 0 were assigned the pseudo-value 1.

Fitness calculations were undertaken using the following formula: Fitness = (read_counts_saline_control / read_count_antibiotic_treated) x (#insertion_sites_antibiotic_treated – #insertion_sites_saline_control). The cluster of orthologous groups were assigned according to the pseudomonas.com database.

## Microscopy

For qualitative and quantitative analysis of shape, size, and fluorescence emission, aggregates were imaged using a Zeiss LSM 980 confocal laser scanning microscope (CLSM) (Zeiss, Germany). The system was running on ZEN 2.1 (Zeiss, Germany). Agar-embedded bacterial aggregates were stained by applying 100 µL of a mixture of 5 µM Syto9 and 20 µM of Propidium iodide (PI) (Live/Dead staining) (Life Technologies, USA) on top of a plug prepared as for CFU determination (Ø 1cm) and cutting a profile section of the agar and placing it on the side of an objective glass, and applying a coverglass on top. The stains were allowed to penetrate the agar for 30 min before imaging. Live/dead stained aggregates were imaged using a 488 nm and a 594 nm laser for excitation with emission filters ranging from 495 – 550 nm and 600 – 640 nm for Syto9 and PI, respectively. Yfp- and Cfp-tagged *P. aeruginosa* strains ^51^ were imaged using a 514 nm and 445 nm laser for excitation with emission filters ranging from 527 – 735 nm and 451 – 517 nm, respectively.

Whole aggregate images were obtained with a 20x #0.4 air objective. Cell-level high-resolution images of aggregates were obtained using a 63x #1.4 oil objective. Images were processed with Imaris 9.5 (Bitplane, Switzerland).

Because the ciprofloxacin concentration used throughout the study results in a 1-2 log10 reduction in CFUs, CLSM is not suitable to compare the effect of ciprofloxacin treatment on the aggregates. Therefore, CFU counts are used to assess and compare the antibiotic response of different mutants.

## Data availability

All data generated or analyzed during this study are included in this published article or its supplementary information files. The DNA sequence data for the study have been deposited in the European Nucleotide Archive (ENA) at EMBL-EBI under accession number PRJEB126817 (https://www.ebi.ac.uk/ena/browser/view/PRJEB126817).

## Supporting information

Supplementary material for Goltermann et al 2026

## Acknowledgments

This work was supported by grants from the Danish Council for Independent Research (grant no. 9040-00023B) and the Novo Nordisk Foundation (NNF22OC0080132 and NNF25OC0107716) to TTN, and the Swiss National Science Foundation (SNSF) (project 310030_192800) and the UZH Global Strategy and Partnerships Funding Scheme to LE.

## Author Contributions

L.G., S.G., K.N.K., L.E. and T.T.N. designed research. L.G., S.G., M.R., S.S., Z.L., S.L. and K.N.K. performed laboratory experiments. T.T.N., S.G. and L.E. contributed reagents/analytic tools. L.G., S.G., M.R., S.S., K.N.K., L.E. and T.T.N. analyzed data. L.G., S.G., M.R., K.N.K., L.E. and T.T.N. wrote the paper.

