## Supplementary material for Goltermann et al 2026 for "Tn-Seq-based identification of genes that play a role in antimicrobial tolerance of *Pseudomonas aeruginosa* aggregates"

#### Supplementary Figures

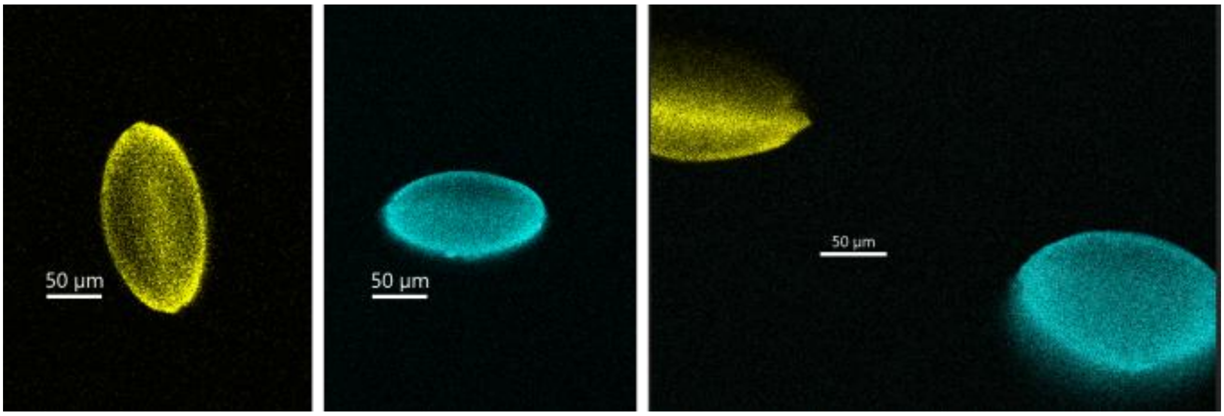


**Figure S1.** Aggregate formation by a mixed PAO1-yfp/PAO1-cfp inoculum. Confocal laser scanning micrographs of 16-hour-old *P. aeruginosa* aggregates originating from a mixed PAO1-yfp/PAO1-cfp inoculum. Twenty-five aggregates were imaged, and all aggregates were composed of bacteria with only one type of fluorescent tag, indicating that they were monoclonal. The images show representative aggregates. The distance of the aggregates was generally so long that only one aggregate was present in each viewing field. However, the image to the right displays a rare case where two aggregates were present in the same viewing field. The size bars correspond to 50 µm.

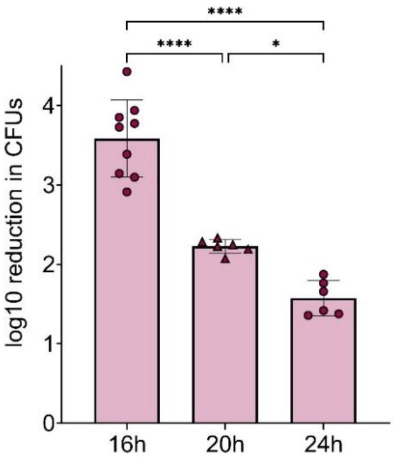


**Figure S2.** Ciprofloxacin treatment of aggregates at different growth stages. Aggregate formation time greatly impacts the effect of ciprofloxacin treatment. Aggregates of 16 h, 20 h or 24 h formation time were treated with 2 µg $mL^{-1}$ ciprofloxacin. The reduction in CFUs after treatment depended on the formation time of the aggregates, with older aggregates being less susceptible. Data was compared by one-way ANOVA, * $p < 0.05$, **** $p < 0.0001$.

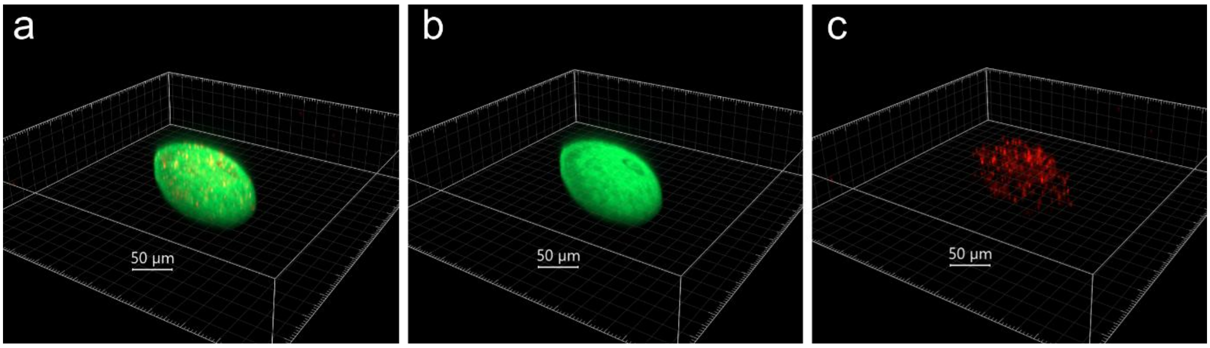


**Figure S3.** Confocal laser scanning micrographs of a Live/Dead stained 16-hour-old *P. aeruginosa* aggregate. Living cells stain green while red color indicates dying bacteria with compromised membranes or extracellular DNA. Z-stacks of the aggregate are shown as 3D presentations with both channels (a) and each of the two channels Cyto9 (b) and PI (c). The size bars correspond to 50 μm.

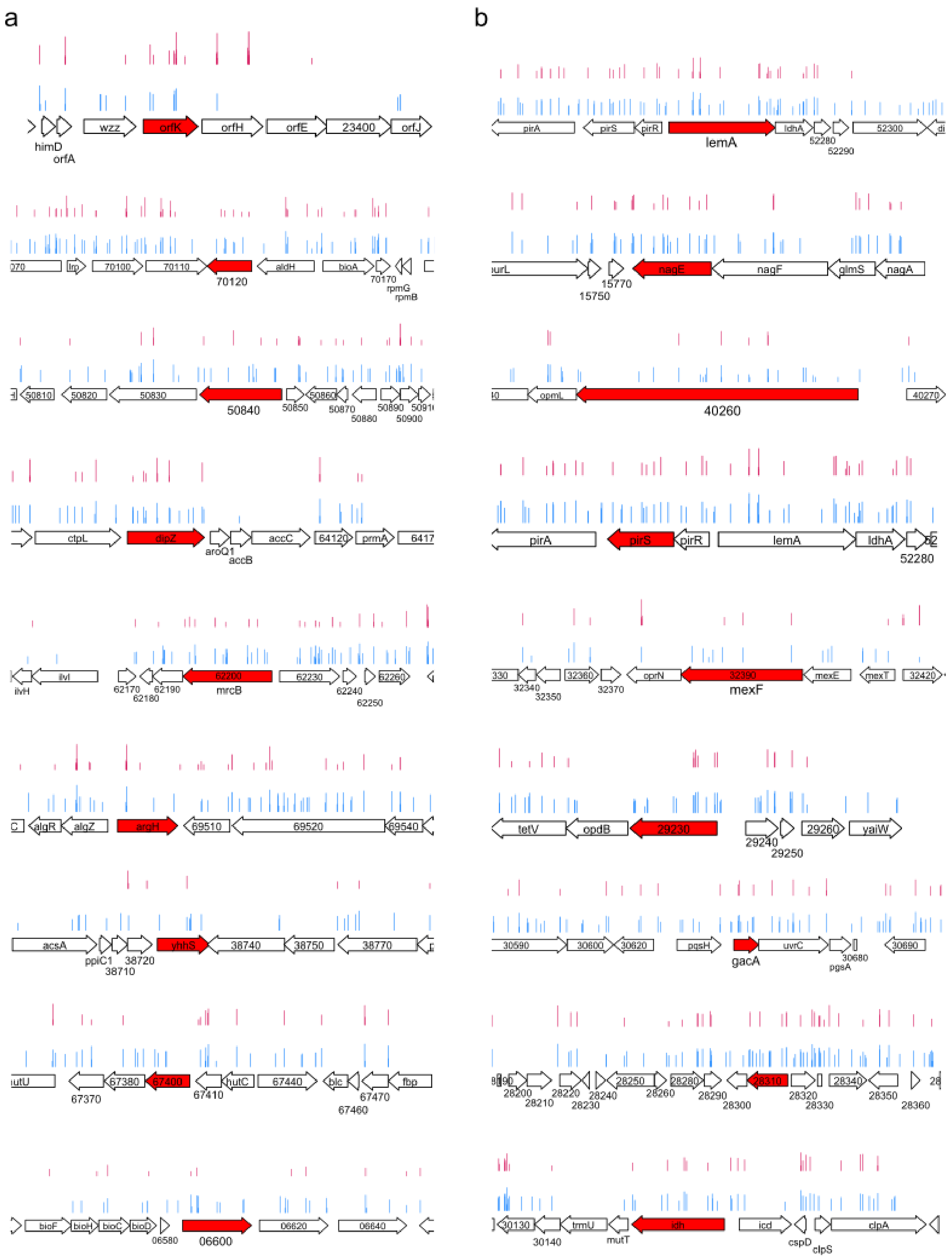


**Figure S4.** Insertion plots. (a) Insertion plots from the tobramycin selection for the top 10 genes identified in the ciprofloxacin selection and (b) insertion plots from the ciprofloxacin selection for the top 10 genes identified in the tobramycin selection.

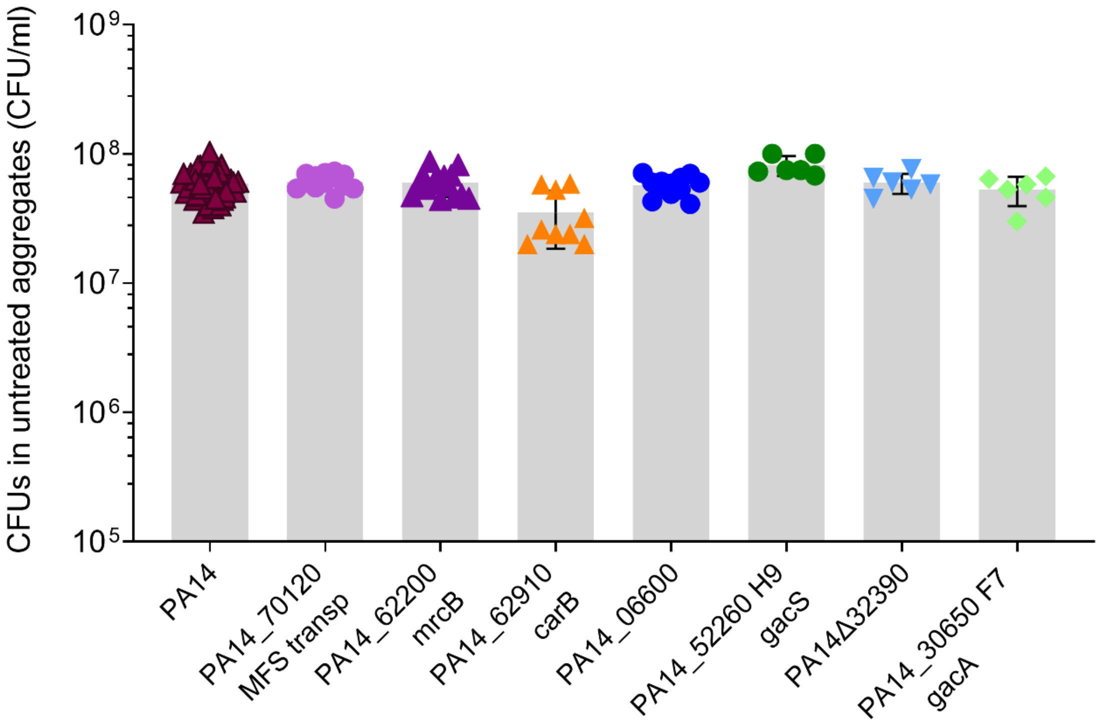


**Figure S5.** Number of viable cells in untreated aggregates. Aggregates were formed for 16 h and treated for 24 h with saline. The number of CFUs is comparable between the wildtype and the transposon mutants. Only the *carB* mutant contained about half the amount of viable cells compared to the wildtype. Means are indicated by bars, and the number of replicates is indicated by the number of dots. Graph shows all combined data points from across all experiments. For statistical analysis, each mutant was compared to the wildtype originating only from the same experiment and data compared using Student's t-test **** $p < 0.0001$.

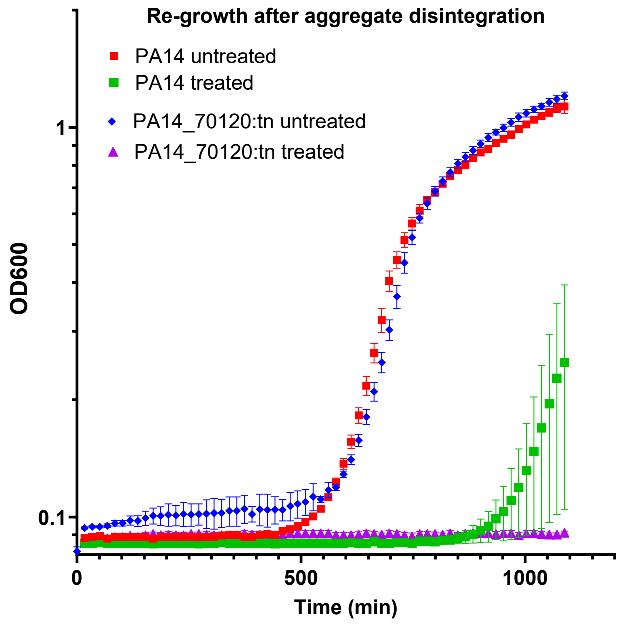


**Figure S6.** Re-growth after ciprofloxacin treatment. Re-growth of bacteria recovered after 24 h ciprofloxacin treatment of 16 h aggregates.

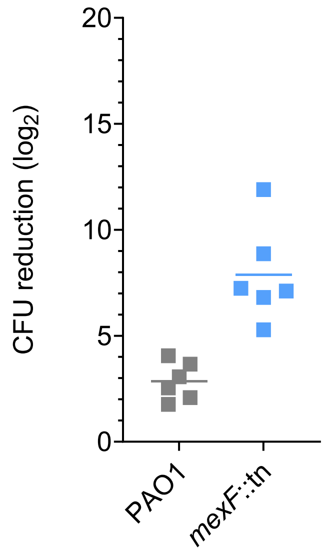


**Figure S7.** Tobramycin tolerance of aggregates formed by the *P. aeruginosa* PAO1 *mexF* transposon mutant and the wild-type PAO1.

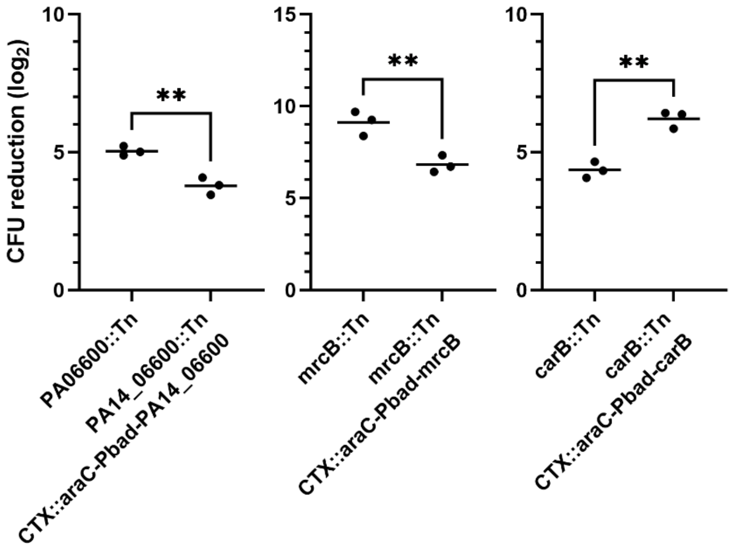


**Figure S8.** Antibiotic tolerance of aggregates formed by the indicated *P. aeruginosa* PA14 transposon mutants and complemented strains. The respective *P. aeruginosa* strains were grown as agar-embedded aggregates for 16 hours, and the aggregates were then treated for 24 hours with ciprofloxacin. Next, the antibiotic was removed by washing and the aggregates were disintegrated, serially diluted and plated for enumeration of viable bacteria (CFUs) and calculation of the log2-reduction of CFUs mediated by the antibiotic treatment compared to saline-treated controls. Arabinose at 0.4% was added to induce $P_{bad}$-controlled genes. The presented data consists of three biological replicates. Means are indicated by lines, and the significance of the difference between the values is assessed by using Student's t-test; ** $p<0.01$.

#### Supplementary tables

##### Table S1. Primers used in this study.

| **Oligonucleotides for Tn-seq** | **Sequence 5'-3'** |
|---|---|
| PAIR_ADAPT_6BC##_HI | ACACTCTTTCCCTACACGACGCTCTTCCGATCTBBBBBBT |
| PAIR_ADAPT_6BC##_LO_5PH | BBBBBBAGATCGGAAGAGCGGTTCAGCAGGAATGCCGAG |
| T23_PAIR_COLLECT | CTTCGGCGCGCCCTAGGGGGATCCTCGGCATTCCTGCTGAACCGCTCTTCCG ATCT |
| T23_SLXA_PAIR_Amp | GGAACTTCGGAATAGGAACTTCTTAGATGTGTATAAGA G |
| SLXA_PAIR_REV_AMP | CAAGCAGAAGACGGCATACGAGATCGGTCTCGGCATTC CTGCTGAACCGCTCTTCCGATCT |
| T23_SEQ_G (Fw) | ATTAGGAACTTCGGAATAGGAACTTCTTAGATGTGTATAAGAGACAG |
| T23_INDEX_1 (Index) | CTAGAGAATAGGAACTTCGGAATAGGAACTTCTTAGATGTGTATAAG |
| PE_READ2_SEQ (Rv) | CGGTCTCGGCATTCCTGCTGAACCGCTCTTCCGATCT |
| **Primers for transposon verification** | **Sequence 5'-3'** |
| PA14_70120 | tcactcggaggcttccag / atgcgttggggaagctatttc |
| PA14_62200 | tcaattcagccagcctcg / atgacgcgtccccg |
| PA14_06600 | atgcctgattacaaggccc / tcagtaaccgagggcg |
| PA14_62910 | cctacaaccgtgaagagttcg / ccgccggtgatgtcg |
| PA14_30650 | gtgattaaggtgctggtggtc / ctagctggcggcatcg |
| PA14_52260 | gtgttcaaggatctcggcatc / tcagagttcgctggagtcg |
| **Primers for k/o construction** | **Sequence 5'-3'** |
| PA14_32390 Up_fw | GGGGACAAGTTTGTA CAA AAA AGC AGG CTC A gaacagatcaagctcccaggcg |
| PA14_32390 Up_rv | cggcgataggaagaaccgatgaatttc gtgcatgcatgattcacgcgc |
| PA14_32390 Down_fw | gaaattcatcggttcttcctatcgccg |
| PA14_32390 Down_rv | GGG GAC CAC TTT GTA CAA GAA AGC TGG GTA caaccaggtcaacccgcgtac |
| PA14_32390 Seq_fw | cagctgacggcggatg |

**Table S2. Results of Tn-Seq.**

This is provided as a separate Excel file.

**Table S3. MIC and growth rate of tested transposon and deletion mutants.**

| | MIC CIP (µg/ml) | MIC TOB (µg/ml) | Growth rate ($h^{-1}$) |
|---|---|---|---|
| **PA14** | 0.125 | 1 | 1.19 (0.11) |
| **70120** | 0.125 | 1 | 1.06 (0.01) |
| **50840** | 0.125 | 1 | 1.19 (0.12) |
| **62200** | 0.125 | 1 | 1.22 (0.11) |
| **06600** | 0.125 | 0.5 | 1.10 (0.08) |
| **62910** | 0.125 | 1 | 0.95* (0.10) |
| **52260 H9** | 0.125 | 1 | 1.15 (0.07) |
| **30650 F7** | 0.125 | 1 | 1.21 (0.05) |
| **Δ32390** | 0.125 | 1 | 1.23 (0.1) |
| **PAO1** | n.d. | 1 | n.d. |
| **PAO1 *mexF*::tn** | n.d. | 1 | n.d. |

* Significantly different from the wildtype according to t-test.

### Supplementary methods

##### Complementation of individual transposon mutants

Arabinose-inducible single-copy chromosomally integrable complementation vectors based on mini-CTX2 (1) were constructed in the following way using NEBuilder HiFi DNA assembly. PCR fragments were generated of the araC-$P_{bad}$ cassette from pJN105 (2) and the genes to be complemented including their native ribosomal binding site just upstream of the respective open reading frame. The two fragments contained overlapping regions allowing their fusion together with *Sma*I-linearized mini-CTX2 resulting in the complementation vectors pCTXaraC-$P_{bad}$-carB, pCTXaraC-$P_{bad}$-mrcB, and pCTXaraC-$P_{bad}$-06600. Primer sequences are available upon request.

The complementation vectors were introduced in the transposon mutants by conjugation using standard triparental mating, thereby creating the complemented strains carB::Tn CTX::araC-$P_{bad}$-carB, mrcB::Tn CTX::araC-$P_{bad}$-mrcB, and PA14_06600::Tn CTX::araC-$P_{bad}$-06600.


1 Newman, J. R. & Fuqua, C. Broad-host-range expression vectors that carry the L-arabinose-inducible Escherichia coli araBAD promoter and the araC regulator. *Gene* **227**, 197-203 (1999). https://doi.org/S0378-1119(98)00601-5
2 Hoang, T. T., Kutchma, A. J., Becher, A. & Schweizer, H. P. Integration-proficient plasmids for Pseudomonas aeruginosa: site-specific integration and use for engineering of reporter and expression strains. *Plasmid* **43**, 59-72 (2000). https://doi.org/10.1006/plas.1999.1441S0147-619X(99)91441-5
